# Leveraging AI to Automate Detection and Quantification of Extrachromosomal DNA (ecDNA) to Decode Drug Responses

**DOI:** 10.1101/2024.10.23.619848

**Authors:** Kohen Goble, Aarav Mehta, Damien Guilbaud, Jacob Fessler, Jingting Chen, William Nenad, Oliver Cope, Darby Cheng, William Dennis, Nithya Gurumurthy, Yue Wang, Kriti Shukla, Elizabeth Brunk

## Abstract

Traditional drug discovery efforts have largely focused on targeting rapid, reversible protein-mediated adaptations to undermine cancer cells’ resistance to therapy. However, cancer cells also exploit DNA-based strategies, typically viewed as slow, irreversible, and unpredictable changes like point mutations or the selection of drug-resistant clones. Contrary to this perception, extrachromosomal DNA (ecDNA) represents a form of DNA alteration that is rapid, reversible, and predictable, playing a crucial role in cancer’s adaptive response. In this study, we present a novel post-processing pipeline for the automated detection and quantification of ecDNA in Fluorescence in situ Hybridization (FISH) images using the Microscopy Image Analyzer (MIA) tool. Our approach is particularly designed to monitor ecDNA dynamics during drug treatment, providing a quantitative framework to understand how ecDNA enables cancer cells to swiftly and reversibly adapt to therapeutic pressure. This pipeline not only offers a valuable resource for researchers aiming to automate ecDNA detection in FISH images but also sheds light on the adaptive mechanisms of ecDNA in response to epigenetic remodeling agents like JQ1.

## Introduction

Cancer cells are masters of adaptation, often using irreversible genomic changes to survive treatment (Labrie et al., 2022). While traditional drug discovery focuses on blocking temporary, protein-level responses, it overlooks the fast-acting genomic responses on extrachromosomal DNA (ecDNA, **Fig 1A**), which are harder to detect and even harder to target. This blind spot leaves 15% of cancer patients without effective therapies (Kim et al., 2020). The urgent challenges are clear: we must generate new technologies to characterize these genomic responses in order to develop new strategies to target their adaptations and deliver better treatment options for patients with cancers that harbor these aberrations.

**Figure 1.**
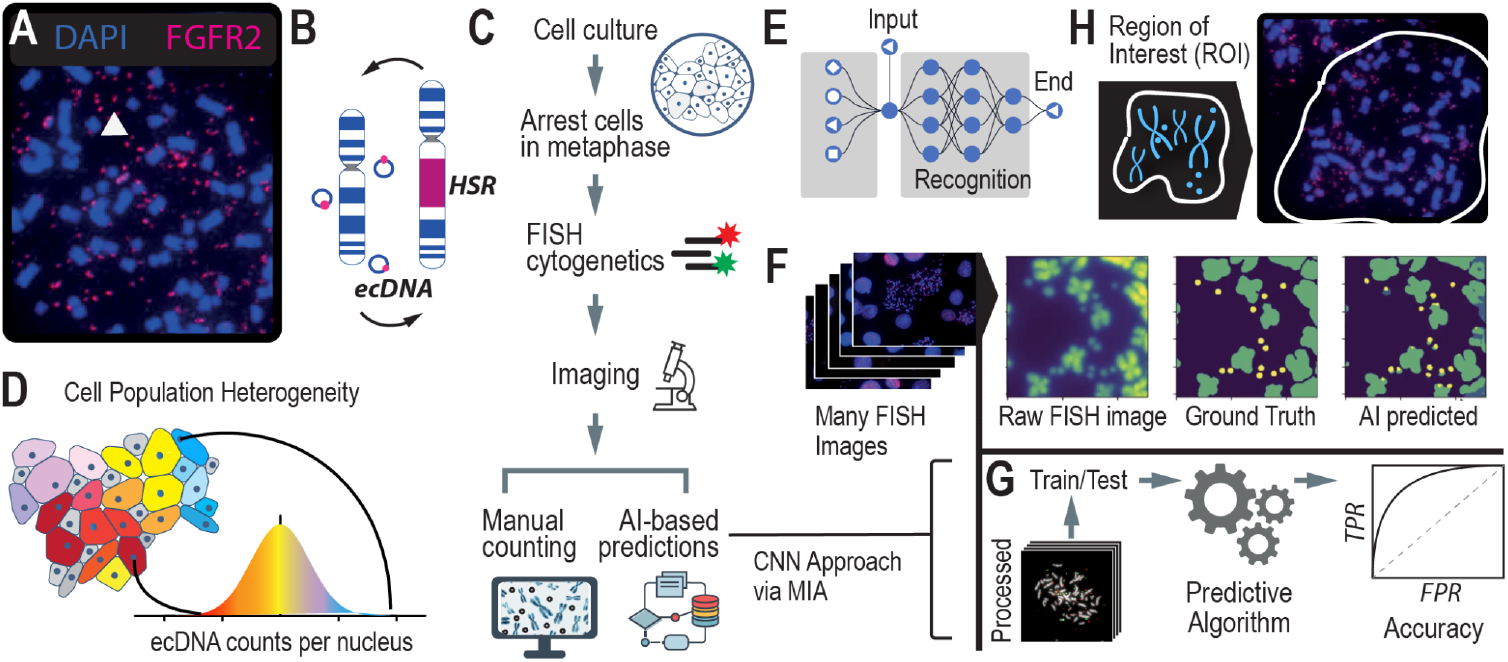
Scaling Annotations of ecDNA in FISH images using AI. **A**. Microscopy methods, such as scanning electron microscopy (SEM) and DNA Fluorescence in situ Hybridization (FISH) are gold standard approaches for visualizing ecDNA. DNA FISH uses fluorescent probes that bind to specific DNA sequences that indicate which regions of the genome are amplified by ecDNA. Multiple species can be observed depending on which fluorescent probe is detected. **B**. Under stress, ecDNAs can re-enter chromosomes to form homogeneous staining regions (HSRs). **C**. FISH is the gold standard method for detecting and analyzing ecDNA in cell nuclei. It consists of several steps which include cell cycle synchronization, fixation, and hybridization. With advances in AI, time-intensive manual labeling and counting of ecDNA is accelerated and can be scaled up. **D**. Asymmetric division of ecDNA molecules into daughter cells during replication and division leads to a heterogeneous population of cells with various ecDNA counts and species. **E**. Convolutional Neural Networks is a computer vision approach that is well suited to image segmentation. **F**. A computer vision approach applied to detecting and counting ecDNAs in FISH images. **G**. Using ground truth (i.e. manually labeled FISH images), we can assess the accuracy of the computer vision method at detecting and counting ecDNAs per cell nucleus.

Next-generation sequencing has revolutionized cancer biology, enabling unprecedented insights into the genetic landscape of cancer. Large-scale consortia such as The Cancer Genome Atlas (TCGA), the Dependency Map (DepMap) (Barretina et al., 2012; Tsherniak et al., 2017), and the Cancer Cell Line Encyclopedia (CCLE) have generated extensive multi-omics datasets, characterizing thousands of cancer genomes and tumor samples. Among these datasets, whole-genome sequencing (WGS), whole-exome sequencing (WES), and ATAC-seq are the most comprehensively covered across samples. Leveraging this data, computational algorithms (Yang et al., 2023)(Deshpande et al., 2019)(Yang et al., 2023) have been developed to detect unique structural variants, including ecDNA and double minute chromosomes (DMs), which frequently appear in cancer cell nuclei. These advancements have fundamentally shifted our understanding of cancer genome structure and karyotype, opening new avenues for research and potential therapeutic interventions.

However, despite their transformative potential, sequencing-based approaches have significant limitations in practice. While they are effective at identifying amplified genes and reconstructing the circular structures of the most dominant ecDNA species, their prediction accuracies typically range between 60-70%(Fessler et al., 2024). This level of accuracy is often insufficient for precise detection and characterization, particularly when detecting ecDNA in cells with fewer copy number counts. Additionally, a major limitation of sequence-based prediction algorithms is that they are not able to distinguish between ecDNA and other genomic elements, such as Homogeneous Staining Regions (HSRs), which are chromosomal regions that can also exhibit gene amplification with “copy-paste-amplify” patterns (**Fig 1B**).

In this way, sequencing alone cannot differentiate between the freely mobile ecDNA, which can independently replicate and segregate during cell division, and HSRs, which are stable and integrated into chromosomes. This inability to distinguish between these structures means that sequencing-based methods may miss crucial aspects of cancer biology, such as the role of ecDNA in driving tumor heterogeneity and drug resistance. Therefore, to accurately detect and characterize ecDNA within cancer cell nuclei, it is essential to integrate additional approaches, such as high-resolution nuclear imaging or functional assays, alongside sequencing data. These complementary methods can provide the necessary context and precision, enabling a more comprehensive understanding of ecDNA’s role in cancer.

Currently, the only methods capable of accurately detecting and quantifying ecDNA, as well as determining its precise location within cell nuclei, are gold-standard cytogenetic imaging techniques. Techniques like Fluorescence in situ Hybridization (FISH) and G-banding karyotyping use fluorescence-based methods to visualize ecDNA within the nucleus (**Fig 1C**). In FISH, cells are typically arrested in metaphase when chromosomes are condensed and spread out, allowing for better visualization of ecDNA. The cell nuclei are stained with fluorescent dyes, such as DAPI, to enhance visibility. When the genetic sequence of ecDNA is known, fluorescent probes can be designed to hybridize with specific regions on the ecDNA, providing unambiguous localization within the nucleus. A standard practice in the field involves generating 20-100 images of individual cell nuclei to capture the heterogeneity in ploidy and ecDNA counts across a cell population (**Fig 1D**). However, the subsequent manual annotation and analysis of these images are time-intensive and low-throughput, limiting the scalability of these image-based approaches. This laborious process underscores the need for more efficient methods that can achieve the same level of accuracy without the associated bottlenecks.

Leveraging AI to automate the detection of ecDNA and HSRs in FISH images represents a transformative shift from traditional, labor-intensive karyotyping methods. By training computer vision models to identify ecDNA in FISH images (**Fig 1E**), we can overcome the historical challenges of scalability, enhancing both accuracy and precision in detection. This approach complements sequence-based prediction methods by providing a robust alternative that directly visualizes ecDNA. Several efforts(Turner et al., 2017; Rajkumar et al., 2019) have been made to automate image-based detection of ecDNA using various approaches, such as thresholding and Convolutional Neural Networks (CNNs, **Fig 1E**). These models employ supervised learning, where they are trained on manually labeled imaging data (i.e., “ground truth” data), with each ecDNA precisely identified and annotated by experts. These annotations supply the model with the exact coordinates of ecDNA in each image, enabling the algorithm to learn the features necessary to predict the presence of ecDNA in non-annotated images (**Fig 1F**). Accuracy of the model is determined by comparing the predicted counts or locations of ecDNA within FISH images to the ground truth data (**Fig 1G**).

Autodetection of ecDNA in FISH images remains a significant challenge due to several factors. FISH images are often noisy, with considerable cell-to-cell variation that complicates traditional image processing techniques. Key issues that reduce prediction accuracy include high noise ratios in pixel intensities, variations in the size and morphology of chromosomes versus ecDNA, and a severe class imbalance where the majority of the image is background or chromosomes. Existing algorithms(Turner et al., 2017; Rajkumar et al., 2019) struggle to generalize across different image types, particularly when transitioning from low-resolution, black-and-white, tiled images to high-resolution images. These models, trained on lower-quality images, are heavily dependent on the specific microscope and image processing conditions, making them non-transferable. As a result, they detect only 30-40% of the ecDNA in our images and suffer from a high false negative rate, especially when ecDNA is close to chromosomes. Moreover, these models are outdated and not publicly available for retraining. There is a critical need for updated, publicly available CNN-based models that can be retrained with higher-resolution images to enhance the accurate detection of ecDNA in FISH images.

Recently, the Microscopy Image Analyzer (MIA) was developed as an end-to-end interface designed to help researchers scale image analysis across large datasets. MIA is unique in its ability to integrate image processing, analysis, and interpretation within a single platform, providing a streamlined approach that significantly reduces the time and effort required for manual annotation. We applied MIA to our specific challenge of auto-detecting ecDNA in FISH images, leveraging its capabilities to create a high-throughput post-processing pipeline (**Fig 1F**). This pipeline dramatically increased the accuracy of ecDNA detection by refining image segmentation, reducing noise, and improving feature recognition. Currently, the field lacks standardized methods for processing and analyzing FISH images, making our contribution particularly valuable for researchers interested in automating ecDNA detection. Additionally, we have applied this approach to pharmacological studies to monitor changes in ecDNA during drug treatment. These experiments require the analysis of numerous images across many conditions, making the ability to scale and autodetect essential for reducing the time and labor involved in such studies.

## Results and Discussion

### Image Processing Pipeline for MIA

The quality of images and annotations plays a crucial role in the success of computer vision models, particularly in tasks such as detecting ecDNA in non-annotated images. High-quality images with precise annotations are essential for the computer to accurately learn and identify the critical features necessary for reliable ecDNA detection. Poor image quality or inaccurate annotations can lead to misrepresentations of features, ultimately undermining the model’s performance and predictive accuracy. In this section, we discuss the comprehensive steps we took to post-process our image database, focusing on rigorous image ranking and quality control (QC) measures. By ensuring that only the highest quality images were used for training and validation, we aimed to optimize the performance of our MIA-based model for ecDNA detection.

#### Pre-processing of FISH Images

##### Accessing Image quality

Image quality can vary due to several technical and biological factors, including the type of cell line used, cell-to-cell variations, post-imaging processing, sample handling, probe potency, and the characteristics of the microscope. A raw image must undergo post-imaging processing to qualify for counting. It is common for any human to over- or under-process an image and incorrectly annotate ecDNAs or HSRs, which generates false negatives and/or positives. Additionally, the biology of the nuclei itself introduces variability. For instance, each image may contain one or more cell nuclei that have burst open, exposing chromosomes and ecDNA. However, the way these nuclei burst on the microscope slide is non-uniform and can occur on slightly different planes, resulting in some nuclei being more in focus than others. Cell-to-cell karyotypic differences after bursting, such as the number and clustering of chromosomes, also greatly fluctuate visibility of ecDNA. This variability makes counting ecDNA within a single nucleus subjective and can lead to inconsistencies in manual annotation. Moreover, the use of DNA probes involves heating samples to accelerate probe hybridization, which can degrade the samples and make them more difficult to image accurately. It is important to create a database of images from which an AI algorithm can train on that are consistent and generalizable. Any overlapping nuclei, debris clouds, or low resolution images decrease the confidence of attribution and negatively impacts the ability of the algorithm to learn correct features for future predictions. While we found that it can be rare to find extremely well resolved and separated bursted nuclei that generate data with high confidence, we tagged and excluded images with lower confidence from training using a systematic grade-based approach.

We ranked images on a scale from 0 to 4 based on several key quality parameters:

###### 1. Resolution and Focus

This parameter assesses the sharpness and clarity of the chromosomes in the image. Images where chromosomes are blurry and indistinguishable, such that the chromosome arms cannot be clearly seen, receive a lower score (0-1). Conversely, images with sharp, well-defined chromosomes receive a higher score (3-4).

###### 2. Uniformity

Uniformity refers to the evenness and consistency of the image. This is evaluated by observing the spatial distribution of the bursted nuclei. If the nuclei are unevenly spread, with overlapping or indistinct regions, the image is marked lower. If the nuclei are evenly distributed with clear boundaries and minimal overlap, the image receives a higher score.

###### 3. Spatial Distinction

This parameter assesses how well the bursted nuclei and chromosomes are separated from surrounding artifacts or debris. If a nucleus is too clustered or overlaps with its surroundings, making it difficult to distinguish individual chromosomes, the image is ranked lower (0-1). Images with well-separated, distinct nuclei that have a clear “splash zone” with no overlap are ranked higher (3-4).

###### 4. Debris and Artifacts

The amount of debris or artifacts in the image is also considered. Images with minimal debris and clean backgrounds are ranked higher, while those with significant debris or distracting artifacts are ranked lower.

Images marked as “0” represent the lowest quality, characterized by poor resolution, lack of uniformity, and high debris, while images marked as “4” are of the highest quality, with clear, well-focused chromosomes, uniform distribution, and minimal artifacts. This ranking system ensures that only the best quality images are used for further analysis and model training.

##### Assessing Annotation quality

Annotation quality in FISH images for counting ecDNA is a complex and time-consuming task, often leading to inconsistencies due to its subjective nature. Different researchers may label ecDNA differently, resulting in variations in the data. Typically, FISH images use DAPI dye to localize ecDNA, which stains all nucleotides but does not differentiate between chromosomal DNA and ecDNA. While using DNA probes that hybridize to specific regions of ecDNA could reduce ambiguity, the exact sequences of ecDNA are not always known. Additionally, these probes are expensive, and there can be heterogeneity in ecDNA sequences within a single sample, leading to incomplete labeling if some ecDNA contain the targeted genes and others do not.

Given these challenges, it is more practical to develop algorithms that predict the location of ecDNA in FISH images using DAPI dye alone. However, different researchers may employ various methods to annotate ecDNA using DAPI, such as adjusting contrast to differentiate between DNA and debris or comparing the general shape and size of potential ecDNA across images. These subjective approaches can lead to significant variation in manual annotations, as there are no clear, standardized criteria for what constitutes an ecDNA. Furthermore, the fact that ecDNA can change shape and size from cell to cell complicates the establishment of consistent annotation rules, making it difficult to generalize across samples.

The annotation of the 3,000 images used in this study was carried out by multiple students over several years. Consequently, the annotation styles, approaches, and quality varied, creating a challenging dataset for machine learning. To address this, our post-processing image ranking system was essential in selecting the most consistent and representative images for training the algorithm. While the model is designed to learn the patterns of ecDNA from the majority of correct annotations, discrepancies in the data—such as errors in labeling of ecDNA that is not visible in the DAPI channel due to its proximity to or overlap with chromosomes—can negatively impact the model’s measured accuracy, making it appear less effective than it actually is.

The images used to train this model were first ranked on a scale from 0 to 4, with 0 indicating images that are highly subjective and difficult to count due to unclear focus, and 4 indicating images that are clearly well-counted and in focus. After an initial ranking, the annotations were reviewed more closely, especially in cases where multiple fluorescent images were merged. Particular attention was given to the DAPI channel, as this is crucial for accurate ecDNA detection. If many annotation markers were found to be skewed, missing, or incorrect, the image received a lower rank. Additionally, notes were taken on potential improvements to enhance image quality for future use.

From the 2,312 images that were reviewed and ranked from previous experiments, about 1,242 images were ranked a 3 or 4 and were chosen to be used for highest quality training data. The table below displays descriptions for each rank and examples in their FISH probe merged with DAPI and grayscale DAPI only with annotations. Images that had little annotation issue were flagged to be perfected and upgraded in ranking.

###### Generating Regions of interest (ROI)

In the context of counting ecDNAs within cell nuclei, generating regions of interest (ROIs, **Fig 1H**) in FISH images is a critical initial step. FISH images typically contain multiple nuclei, some of which may be intact while others are bursted, revealing the underlying chromosomes and ecDNAs. These images can be complex, with overlapping structures and varying degrees of focus, making it challenging to accurately count ecDNAs without precise guidance. By defining ROIs, researchers can direct the algorithm to focus specifically on the areas of interest—namely, the individual nuclei or their remnants—thereby filtering out irrelevant parts of the image. This targeted approach significantly enhances the accuracy of ecDNA counts, ensuring that the algorithm is analyzing only the relevant nuclear material and not extraneous background or overlapping cells. By clearly delineating ROIs, the post-processing steps are streamlined, leading to more reliable and reproducible results in the study of ecDNA within cancer cells.

With the selected set of high quality images, it was essential to address the presence of debris, as well as chromosomes and ecDNA that had drifted from other nuclei, particularly at the edges of the images. These extraneous elements were not originally annotated and needed to be removed to focus on the relevant data. To achieve this, a lasso tool was employed to draw a region of interest (ROI) that encapsulated all annotated ecDNA while minimizing the inclusion of non-annotated material. The resulting ROIs were saved as binary masks. During this process, image contrast was significantly enhanced to differentiate and either include or exclude faintly visible features. For nuclei that were close together but clearly separated, each was carefully reviewed, with distinct cells isolated into their own ROIs when possible or otherwise removed from the dataset. This approach resulted in a variety of ROI shapes, ranging from tight circles to elongated ovals and irregular polygons, effectively tracing around unburst nuclei and isolated debris clouds.

As previously mentioned, subjectivity can arise when interpreting the boundaries of a cell’s bursted nuclei radius and distinguishing between circularized debris and ecDNA, leading to inconsistencies in annotations between different individuals. Defining ROIs not only provides the algorithm with a focused area for counting and segmentation but also reduces the influence of these subjective interpretations. By constraining the analysis to clearly defined areas, we minimize the need for multiple revisions of the boundary during post-prediction review. Additionally, this approach facilitates a more consistent and reliable evaluation of ground truth image annotations, ensuring that they can be accurately reviewed and audited within a well-defined context.

###### Expanding Image Annotations

Our images were annotated by placing a dot on a single pixel to mark the location of ecDNA. However, this dot may not always align perfectly with the pixel of highest intensity for that ecDNA, which can impact the quality of annotations and subsequently affect the performance of machine learning algorithms. Recognizing the importance of precise annotations in training an algorithm, we developed a method to improve these annotations. We created a mask around each annotated pixel, shaped like a diamond, to include nearby pixels surrounding the ecDNA mark. This approach provided a more comprehensive representation of the ecDNA, allowing the algorithm to better identify and classify ecDNA by capturing a broader context of the signal. This enhancement was crucial in improving the accuracy and robustness of our machine learning model.

#### Evaluation Datasets: Human Error, Ceiling Performance, and Full Model Benchmarking

##### Human Error

To better understand the amount of human error involved in annotating FISH images, we conducted an experiment where we selected a subset of FISH images. Three different individuals were asked to annotate this subset independently, and one individual annotated the same subset in three separate replicates. Our findings revealed that across the three individuals, there was a 8% +/-1% error in the annotations, indicating the presence of human error. This analysis suggests that any algorithm trained on such data is inherently limited by this level of technical error, implying that the maximum achievable accuracy for the algorithm is constrained by the variability in human annotations. This establishes a benchmark for understanding the limitations of our model’s performance and provides a realistic expectation for its accuracy (**Fig 2A**).

**Figure 2.**
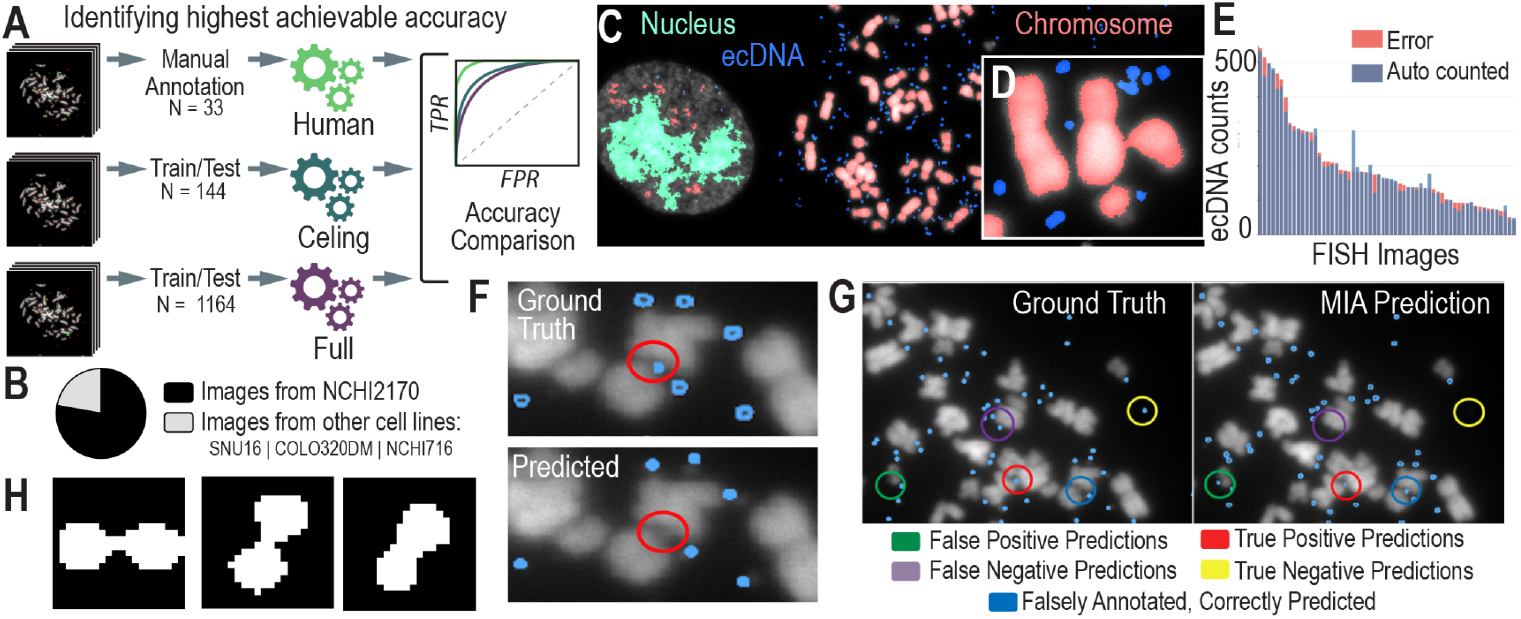
AI Models and Assessment of their Accuracies. **A**. We created different models to test the highest possible accuracy that we could expect our model to reach. **B**. The breakdown of images used to train the Convolutional Neural Networks (CNNs) by cell line model system. **C**. CNN-predicted chromosomes, ecDNA, and in-tact nuclei from a FISH image. **D**. A zoomed-in image of ecDNA near chromosomes, highlighting the challenges in detecting individual ecDNAs when they cluster close together. **E**. Accuracy metrics for the full (n=1164) model, showing the distribution of error (the difference in ecDNAs predicted versus manually counted) across images. **F**. Existing algorithms struggle to predict ecDNAs in close proximity to chromosomes. **G**. Annotated results from MIA, showing instances of true positives/negatives and false positives/negatives as well as cases where MIA finds ecDNA that were not correctly annotated.

##### Ceiling Performance

To determine the highest achievable accuracy of MIA, we focused on a subset of hand-picked, top-quality FISH images that were most amenable to automated AI learning. We selected 144 FISH images that ranked highest according to our stringent image quality criteria, ensuring they were clear, well-focused, and exhibited minimal noise or overlap. By training MIA on this cherry-picked dataset, we aimed to create a “ceiling model” that represents the maximum potential accuracy of ecDNA detection under ideal conditions. While we anticipate that this model will demonstrate superior performance, we also recognize that it may not be generalizable to more diverse or lower-quality images. Nonetheless, this approach provides valuable insight into the upper limits of algorithmic accuracy, establishing a benchmark for what can be achieved in optimal scenarios.

##### Full Model

To develop a robust and generalizable model, we created a full model dataset by selecting images with a ranking of 3 or higher, ensuring a higher standard of image quality for training. This dataset comprises 1164 images, with 78% derived from our primary model system, NCIH2170 cells, while the remaining 22% come from a diverse set of cell line model systems that harbor ecDNA, including NCIH716, SNU16, and COLO320 (**Fig 2B**). By incorporating a variety of cell lines, this dataset aims to strike a balance between achieving high accuracy and maintaining broad applicability across different cellular contexts. The full model represents our comprehensive effort to train an algorithm capable of accurately detecting ecDNA across a range of image qualities and biological conditions.

#### Segmentation

Segmentation in computer vision is the process of dividing an image into distinct segments or regions, where each segment corresponds to different objects or parts within the image. The primary objective of segmentation is to simplify or transform the representation of an image into something more meaningful and easier to analyze, making it a critical step for tasks like ecDNA recognition.

Semantic segmentation is particularly effective for identifying and analyzing ecDNA within cell nuclei in FISH (Fluorescence in situ Hybridization) images, as demonstrated in previous approaches (Rajkumar et al., 2019). By classifying each pixel in an image into predefined categories—such as “ecDNA,” “chromosome,” “unbursted nuclei,” or “background”—semantic segmentation treats all objects of the same category as a single class (**Fig 2C**). While it doesn’t distinguish between different instances of the same object, it excels in detecting and segmenting ecDNA within the complex cellular environment (**Figure 2D**). This approach overcomes challenges posed by overlapping structures and varying signal intensities in FISH images, enabling more accurate analysis of ecDNA.

Microscopy Image Analyzer (MIA) builds on these principles by incorporating various image processing techniques, including semantic segmentation, depending on the specific model or task. MIA is designed for flexibility and adaptability across different types of microscopy images, including those used for ecDNA detection in FISH images. While it can employ semantic segmentation similar to previous approaches, MIA primarily focuses on streamlining the analysis of large-scale image datasets through object detection, classification, and region-of-interest (ROI) selection (**Fig 1H**). By applying techniques like thresholding, edge detection, and ROI selection, MIA enables researchers to identify and quantify features of interest—such as ecDNA—in a high-throughput, efficient manner.

##### Accuracy and Performance

We began by testing the accuracy and performance of our ceiling model. Accuracy was assessed by comparing the total ecDNA count in each image within the validation set against the annotated ground truth. This metric guided the initial iterations of model improvement. The first training sets consisted of approximately 144 ultra-high-confidence, hand-picked images. When evaluated against a validation set of 45 selected images, the model exhibited an absolute difference in total counts of around 8% compared to the ground truth. While this level of accuracy was unexpectedly high, it likely represents the upper limit of achievable performance. In subsequent iterations, models were trained on a larger dataset comprising 1164 images, which were refined by selecting ROIs and filtered to include only images with ranks of 3 and 4. After incorporating contour detection (discussed below) to address conjoined segmentations, the error rate for the resulting models, evaluated on a new set of 55 higher quality images (33 ranked 3+), was 7.62% (**Fig 2E**).

After achieving accuracy levels comparable to human annotations, we introduced an additional precision metric to further evaluate model performance. This metric involved comparing the predicted segmentation masks generated by the model to the ground truth masks (created using a pixel-patched approach to address class imbalance). Specifically, we assessed whether each unique object in the ground truth had at least one pixel overlap with a predicted mask, considering these overlapping objects as correctly identified. However, we refined the process to ensure that each object in the ground truth was exclusively matched to a single predicted object, preventing multiple predictions from being counted as overlaps with the same ground truth object. This approach helped us identify and balance false positives and negatives that might otherwise have been missed, thereby improving the overall quality of the training set.

##### Comparison to Existing Approaches

Previous approaches to automating ecDNA detection, such as ecDetect and ecSeg(Turner et al., 2017; Rajkumar et al., 2019), have made significant strides in addressing the challenges of accurately counting and localizing ecDNA within cell nuclei. ecDetect primarily relies on size thresholding to distinguish ecDNA from other nuclear components, applying a filter to isolate the region of interest (ROI) before analysis. However, this method can struggle with noise and closely clustered ecDNA, leading to potential inaccuracies in annotation and resolution. ecSeg, on the other hand, utilizes a U-Net architecture with a ResNet50 backbone and focuses on segmenting images into patches for analysis. While effective, ecSeg was trained on a relatively small dataset of 483 annotated images, which may limit its generalizability and resolution at the chromosomal level.

Using the Microscopy Image Analyzer (MIA) tool builds on these previous methods by incorporating advanced image post-processing techniques that enhance annotation quality and improve resolution. Trained on nearly 1,300 high-quality images, MIA benefits from a larger and more diverse training dataset and introduces a refined segmentation process that accurately identifies and separates closely clustered ecDNA, resolving them even when colocalized around or between chromosomes (**Fig 2F**). Unlike ecDetect, MIA’s method does not rely heavily on size thresholding, allowing it to maintain high accuracy without filtering out subtle ecDNA signals.

Moreover, MIA’s ability to classify chromosomes within the segmented ROIs provides an additional layer of precision, which is crucial for distinguishing between ecDNA and chromosomal artifacts. This capability directly impacts the overall accuracy of ecDNA detection, as it reduces false positives and negatives that arise from misclassification in complex cellular environments. By comparing the performance of MIA with ecDetect and ecSeg (**Table 1, Sections A and B, and Supplementary Table 1**), we demonstrate significant improvements in both the accuracy of ecDNA identification and the ability to handle unannotated images.

**Table 1.** Accuracy metrics across all models. In Section A, MIA is the CNN-based model and we assess accuracy for the Ceiling, 2170 (with 3+ or higher ranked images from only NCIH2170 cells), and Full models. The Full model was split twice to evaluate the differences between training on different images in the dataset. In Section B, ecSeg is the CNN-based model used to assess all but the Ceiling model. In Section C, we use the Full MIA model to predict ecDNA counts in FISH data from control NCIH2170 cells and cells that have been treated with JQ1. We also assess the Full ecSeg model to predict ecDNA counts in the same set of images. In Section D, we use the Full MIA and Full ecSeg models to predict ecDNA counts in other cell lines treated with JQ1 to evaluate the generalizability of these models to other cell lines.

| Model | Pred count | Actual count | Accuracy | Precision | Recall | Matthew | F1 |
| --- | --- | --- | --- | --- | --- | --- | --- |
| **A. MIA Model Metrics** |  |  |  |  |  |  |  |
| Ceiling | 203.016 | 193.778 | 0.83 | 0.842 | 0.877 | 0.653 | 0.857 |
| 2170 (3+ images) | 212.184 | 215.868 | 0.796 | 0.835 | 0.817 | 0.587 | 0.824 |
| Full Model | 157.728 | 214.927 | 0.637 | 0.724 | 0.561 | 0.305 | 0.621 |
| CV Full Model | 132.668 | 180.703 | 0.621 | 0.709 | 0.538 | 0.279 | 0.597 |
| **B. EcSeg Model Metrics** |  |  |  |  |  |  |  |
| 2170 (3+ images) | 191.197 | 215.868 | 0.656 | 0.712 | 0.641 | 0.324 | 0.667 |
| Full Model | 79.31 | 214.927 | 0.424 | 0.441 | 0.199 | -0.058 | 0.278 |
| CV Full Model | 78.81 | 180.703 | 0.438 | 0.435 | 0.237 | -0.05 | 0.315 |
| **C. Drug Response Predictions** |  |  |  |  |  |  |  |
| 2170 control (MIA) | 284.532 | 326.675 | 0.719 | 0.796 | 0.696 | 0.444 | 0.74 |
| 2170 JQ1 (MIA) | 100.758 | 165.379 | 0.603 | 0.777 | 0.47 | 0.281 | 0.579 |
| 2170 control (ecSeg) | 97.286 | 326.675 | 0.413 | 0.505 | 0.164 | -0.048 | 0.242 |
| 2170 JQ1 (ecSeg) | 40.121 | 165.379 | 0.375 | 0.399 | 0.101 | -0.147 | 0.166 |
| **D. Drug Response Predictions in other cell lines** |  |  |  |  |  |  |  |
| COLO320 JQ1 (MIA) | 35.828 | 45.969 | 0.544 | 0.62 | 0.481 | 0.149 | 0.535 |
| COLO320 JQ1 (ecSeg) | 10.328 | 45.969 | 0.347 | 0.338 | 0.078 | -0.19 | 0.184 |
| SNU16 JQ1 (MIA) | 142.08 | 260.3 | 0.658 | 0.738 | 0.522 | 0.382 | 0.571 |
| SNU16 JQ1 (ecSeg) | 18.8 | 260.3 | 0.335 | 0.119 | 0.013 | -0.213 | 0.039 |

#### Post-processing analyses

##### Contour detection

Contour detection is a technique used in image processing and computer vision to identify and delineate the boundaries or edges of objects within an image. A contour represents a curve that connects all the continuous points along a boundary that share the same color or intensity. In the context of image segmentation, contour detection is crucial for accurately separating and defining the shapes and structures of different objects within an image, such as nuclei, chromosomes, or ecDNA.

When dealing with conjoined segmentations—instances where multiple objects (e.g., closely clustered ecDNA) are touching or overlapping (**Fig 2H**), contour detection helps to identify the precise boundaries of each individual object. This allows for the separation of these conjoined objects into distinct segments, thereby improving the accuracy of the model in distinguishing between different ecDNA entities or other features in the image.

To enhance the accuracy of ecDNA detection, we applied contour detection as a critical post-processing step. By accurately delineating the boundaries of individual ecDNA within these complex regions, we were able to significantly improve the model’s ability to distinguish between different ecDNA entities. The improvement after using contour detection underscores the importance of advanced post-processing techniques in refining the performance of computer vision models for complex biological image analysis.

### Merging Data from Other Fluorescent Channels

Once MIA was executed to predict ecDNA in unannotated images, our next step was to identify the specific genes located on these predicted ecDNA, enabling us to count and differentiate between different ecDNA species and assess oncogene amplification levels. In our case, we labeled two oncogenes known to localize to ecDNA in each cell line: ERBB2 and MYC for NCIH2170 cells, and FGFR2 and MYC for SNU16 and NCIH716 cells. Beyond using DAPI fluorescence, which provides a general overview of nucleic acids in our images, we employed FISH DNA probes that bind directly to genomic regions within each of these genes. To accurately map the predicted ecDNA locations with the actual locations identified through other channels (e.g., FITC and Texas Red), we developed a post-processing script. This script enabled the alignment of MIA-predicted ecDNA with the fluorescent signals from the gene-specific probes. When comparing the results of this script to human annotations, we found that the accuracy ranged within 97% ± 2%(**SI Figure 1-3 and SI Table 2**), demonstrating the effectiveness of our approach (See **Supplementary Methods** for more details).

### Applications in Systems Pharmacology: Changes in ecDNA During Drug Treatment

Approximately 15% of cancers harbor ecDNA(Kim et al., 2020), yet research into the critical response mechanisms involving ecDNA remains limited, and only a few drugs have been thoroughly explored. Identifying treatments that specifically target cells with ecDNA presents a significant challenge. Without a rational, systematic approach, drug selection can become a time-consuming and resource-intensive process of trial and error. To address this challenge, we developed CytoCellDB(Fessler et al., 2024), a comprehensive global data resource designed to streamline drug response evaluations. This platform allowed us to systematically assess drug efficacy across 139 cell lines harboring ecDNA and over 400 cell lines without DMs, providing a robust framework for efficiently identifying impactful treatments. Through this approach, we also identified a cell line whose ecDNA was not yet reported, that co-amplifies ERBB2 and MYC on the same ecDNA amplicon, highlighting the potential of CytoCellDB to uncover new insights in cancer research.

Co-amplification of oncogenes on ecDNAs presents unique opportunities for diversification of gene and gene dosage combinations across cells in a population. As demonstrated by **Fig 3A**, there are three identified “species” of unique ecDNA and one unidentified species. These different ecDNA species localize different genes or gene combinations on the same fragment of ecDNA, “mix and matching” genes that are not commonly that close to one another in genomic space. For example, the gene MYC is encoded in chromosome 8 and the gene ERBB2 is encoded in chromosome 17. Suddenly, on ecDNA they become neighbors in the ecDNA species that co-localize these genes on the same amplicon. In NCIH2170’s case, we see ecDNA that only localizes MYC or ERBB2 separately, and those that co-amplify these genes. We also see an “unknown” entity that amplifies genes other than MYC or ERBB2. These different species generate far more population genetic diversity than having one species alone. Increasing genetic population heterogeneity may be a strategy of these cancer cells to diversify their populations to increase the likelihood of generating advantageous traits to overcome drug resistance or changing environments, effectively “hedging their bets” (**Fig 3B**).

**Figure 3.**
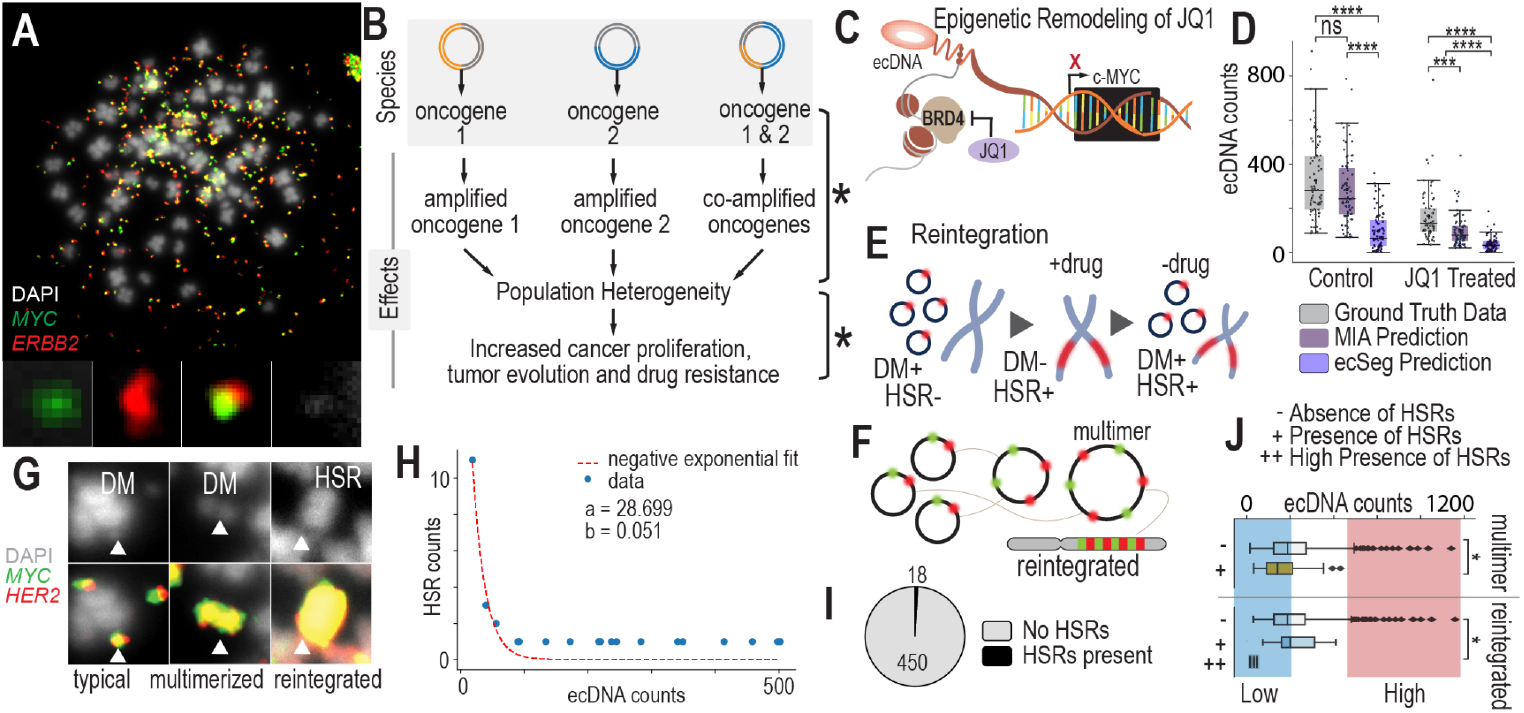
Scaling Data Analytics to Probe ecDNA-Mediated Drug Response Mechanisms. **A**. FISH image with dual probe labeling of NCIH2170 cells that have four or more ecDNA species present in most of their cell nuclei. **B**. Having multiple species of ecDNA present across cell nuclei can generate widespread cell-to-cell heterogeneity. As ecDNA segregate unevenly into daughter cells, each of these species will segregate unevenly. This generates extreme population genetic heterogeneity in terms of copy number variation differences across cells. This heterogeneity could lead to accelerated adaptation and drug resistance. **C**. JQ1 is a molecule that was recently discovered to impact ecDNA higher-order clustering. It inhibits BRD4, impacting MYC transcriptional activity. Because MYC is often amplified on ecDNA, JQ1 may globally influence cells that harbor ecDNA-based MYC amplications. **D**. Experimental results and AI-predicted ecDNA counts for MIA and ecSeg comparing ecDNA counts in control (DMSO treated) and drug treated (JQ1) NCIH2170 cells. **E**. Under treatment, cells eliminate their ecDNA rapidly and reintegrate them into chromosomes, forming homogeneous staining regions (HSRs). **F**. Drug treatment increases the number of multimerized ecDNAs that are seen in FISH images, which may be a precursor to chromosomal reintegration. **G**. FISH data showing different structural views of ecDNA during drug treatment, including double minutes, multimerized ecDNA or “hubs” and HSRs. **H**. A model predicting ecDNA reintegration rate during drug treatment. **I**. The fraction of cells that were observed to have HSRs during FISH image analysis. **J**. Cells with lower ecDNA counts are more likely to be observed to have HSRs compared to cells with high ecDNA levels. Similarly, cells with lower ecDNA levels are more likely to have multimerized structures compared to cells with high levels of ecDNA.

Currently, we lack a systematic method to quantify changes in ecDNA across different therapies. Previous studies that have studied the effects of drugs on ecDNA have been fragmented and inconsistent, with significant variations in cell lines, drugs, dosages, and time points. This lack of standardization has led to an incomplete understanding of the dynamic processes involved in ecDNA drug response. Most research to date has primarily focused on the effects of chemotherapeutic agents (Haque et al., 2001),(Storlazzi et al., 2010),(Kitajima et al., 2001),(van Leen et al., 2022), such as methotrexate, hydroxyurea, cisplatin, etoposide, doxorubicin, and vincristine, on DM reintegration. Interestingly, tyrosine kinase inhibitors (TKIs) have been shown to induce reversible reintegration of DMs (Nathanson et al., 2014), highlighting the complexity of the interactions between drugs, ecDNA and cellular adaptation strategies.

Recent studies suggest that epigenetic remodeling agents, such as JQ1, may exert unique effects on cell lines harboring ecDNA. JQ1 and other epigenetic modulators alter chromatin accessibility, effectively modifying the ‘openness’ or ‘closed-ness’ of DNA regions. Given JQ1’s known interactions with ecDNA, this discovery presents an exciting new avenue for drug development. JQ1 specifically targets BRD4(Zhou et al., 2020), a protein that enhances MYC transcription and tethers DMs within transcriptional hubs(Wu et al., 2019; Hung et al., 2021). Inhibiting BRD4 not only disrupts DM-specific transcription(Wu et al., 2019) but also impacts the broader transcriptional landscape (**Fig 3C**). These alterations can significantly influence downstream processes like gene transcription by regulating which genes are accessible. An unintended consequence of such epigenetic remodeling might be the disruption of ecDNA hubs, potentially undermining a critical adaptive mechanism in cancer cells harboring DMs.

As a pharmacology application, we leverage our AI-accelerated image annotation platform, using the Microscopy Image Analyzer (MIA), to study changes in ecDNA counts before and after drug treatment across three distinct cancer cell lines: a lung cancer cell line (NCIH2170), a gastric cancer cell line (SNU16), and a colorectal cancer cell line (NCIH716). Our goal is to apply our image processing pipeline and MIA to predict ecDNA counts in images from both control and drug-treated samples, with a focus on assessing the impact of JQ1 treatment on ecDNA counts after 24 hours. This approach aims to provide deeper insights into the effects of JQ1 on ecDNA dynamics and its potential role in disrupting critical adaptive mechanisms in cancer cells.

#### Model Performance for Predicting ecDNA Counts in JQ1-Treated Samples

The MIA model and the ecSeg model were evaluated for their ability to predict ecDNA counts in both control and JQ1-treated samples. Our objective was to determine how accurately each model could quantify the reduction in ecDNA levels induced by JQ1 treatment, a BET inhibitor known to impact chromatin accessibility and gene expression.

The MIA model achieved reasonable accuracy in predicting ecDNA counts across both control and treated conditions. For control samples, the model obtained an accuracy of 0.719, with precision and recall values of 0.796 and 0.696, respectively, resulting in an F1 score of 0.74. However, when applied to JQ1-treated samples, the model’s performance declined, reflecting the biological complexity of drug-induced changes. In treated samples, the MIA model reported an accuracy of 0.603, with precision at 0.777 and recall at 0.47, yielding an F1 score of 0.579. The drop in recall suggests that the model undercounted ecDNA, possibly due to the altered ecDNA patterns post-treatment.

The ecSeg model exhibited lower overall performance compared to the MIA model. For control samples, it achieved an accuracy of 0.413 with a precision of 0.505 and a recall of 0.164, indicating difficulty in detecting the full ecDNA population. In JQ1-treated samples, the ecSeg model’s performance further declined, with an accuracy of 0.375 and a precision of 0.399, while recall dropped to 0.101, resulting in a negative Matthew’s correlation coefficient (MCC) of −0.147. These metrics highlight the ecSeg model’s limitations in adapting to ecDNA changes following drug treatment.

#### Model Comparison and Implications

The performance of both models underscores the challenges in predicting ecDNA counts under drug treatment conditions. While the MIA model demonstrated moderate generalizability, achieving reasonable performance across control and treated samples, the ecSeg model struggled with both control and JQ1-treated datasets. The higher precision but lower recall in JQ1-treated samples for both models suggests that while some ecDNA features were accurately identified, a significant portion of ecDNA was missed, possibly reflecting altered ecDNA morphology or reintegration patterns in response to the drug.

These findings indicate that while our models, especially MIA, offer potential for predicting ecDNA behavior, further model refinement is needed to improve recall and robustness under varying biological conditions. The results also demonstrate the need to incorporate additional training data reflecting post-treatment changes to better capture the dynamics of ecDNA in response to drugs like JQ1.

(**Table 1, Sections A and B**)

#### Cells Rapidly Eliminate ecDNA After 24 Hours of JQ1 Treatment

Comparing ecDNA counts before and after 24 hours of JQ1 treatment, we observe that cells rapidly eliminate their ecDNAs. In NCIH2170 cells, the mean ecDNA count in untreated wild-type cells is 330 ± 167, while after JQ1 treatment, the mean ecDNA count drops to 220 ± 132 (**Fig 3D**). Similarly, in COLO320 cells, the mean ecDNA count decreases from 50 ± 37 in wild-type cells to 45 ± 30 post-treatment. In SNU16 cells, the mean ecDNA count shifts from 370 ± 203 in wild-type cells to 298 ± 156 following JQ1 exposure. Using a Wilcoxon rank-sum test, we find that the difference in ecDNA counts between untreated and JQ1-treated populations is highly significant across all three cell lines. This suggests that JQ1 induces a similar drug response mechanism as other chemotherapeutic agents, where cells rapidly eliminate their ecDNA, possibly through the selection of less sensitive clones, cells with fewer ecDNAs, or active elimination processes.

#### Cells Reintegrate ecDNAs into Chromosomal Homogenous Staining Regions (HSRs)

During drug treatment, cells reintegrate ecDNA into chromosomes, forming HSRs, or regions of highly amplified genes within chromosomes (**Fig 3E**). In this way, ecDNA, which is extrachromosomal and independent of chromosomes, shapeshifts into chromosomal DNA. As a field, we still do not definitively understand how this phenomena increases cell fitness or survival during drug treatment. The overarching opinion is that cells “hide” and “store” their ecDNA in HSRs when it will serve them better in future conditions. Furthermore, ecDNAs form higher-order structures in which it appears that they multimerize and combine into larger ecDNA entities (**Fig 3F**). Multimerization may occur prior to reintegration; however, we currently lack sufficient evidence to confirm this process definitively (**Fig 3G**).

Comparing the number of HSR counts in NCIH2170 cells before and after JQ1 treatment, we find that cells reintegrate ecDNAs at a rate of approximately 5% during drug exposure (**Fig 3H**). In this cell line, no HSRs are observed in control cells that contain the same set of genes as the ecDNAs, indicating that any HSR detected in the drug-treated samples containing MYC or ERBB2 represents a reintegrated ecDNA. After 24 hours of drug treatment, we observe 18 reintegration events across 450 cells (**Fig 3I**), confirming that ecDNA reintegration occurs as part of the cellular response to the drug. *ecDNA Reintegration Mostly Occurs in Cells with Fewer ecDNA Counts*.

Reintegration events are more likely to occur in cells that have lower numbers of ecDNA. Analysis of over 600 FISH images from NCIH2170 cells showed that DMs reintegrate into HSRs more frequently in cells with fewer DMs (p=0.027, chi-squared test), consistent with findings from previous studies(Haque et al., 2001) (**Fig 3J**). One possible explanation for this observation is that cells with fewer ecDNAs may face lower cellular allocation costs, allowing them to invest more resources into processes such as reintegration. Alternatively, these cells may have different cell cycle distributions, potentially spending more time in specific phases that favor reintegration events. This could enable a higher frequency of HSR formation in cells with fewer ecDNAs, offering them a survival advantage under drug treatment conditions.

### Model Limitations

While the Microscopy Image Analyzer (MIA) offers significant advantages in automating the detection of ecDNA in FISH images, it does have certain limitations that users should be aware of. One key limitation is the inability to extract data such as output curves directly from the software, which can hinder detailed post-analysis and further processing. Additionally, MIA has substantial memory requirements, which can be a constraint when working with large datasets or on machines with limited resources. The software is also restricted to a graphical user interface (GUI) and lacks command-line functionality, preventing it from running in a “headless” mode—this limits automation and integration into larger, script-based workflows. Moreover, MIA does not allow users to pull data directly from a master folder, requiring manual file management and input, which can be cumbersome for large-scale analyses. These limitations highlight areas for potential improvement in future versions of the software to enhance its utility and flexibility in advanced research applications.

A detailed guide on how to prepare, rank, and analyze images using the MIA with FISH images is provided in the Supplementary Information. This documentation includes step-by-step instructions on image preparation, ranking criteria, and running MIA for optimal results.

## Conclusions

In this study, we successfully automated the detection of extrachromosomal DNA (ecDNA) using the Microscopy Image Analyzer (MIA), providing a robust and accurate framework for analyzing FISH images. Through the development and application of our custom pipeline, we enhanced MIA’s capabilities specifically for ecDNA detection and quantification, establishing it as a powerful tool for investigating the effects of pharmacological treatments on cancer cells.

Our approach has yielded new insights into cellular responses during JQ1 treatment, revealing significant shifts in ecDNA counts, patterns of ecDNA reintegration into chromosomes, and the tendency of certain cells within the population to favor ecDNA reintegration. These findings underscore the potential of MIA-based analysis to advance our understanding of how cancer cells adapt to therapeutic pressures, offering valuable perspectives for drug discovery and cancer treatment strategies.

Additionally, we observed promising signs of generalizability. Models trained primarily on images from one cell line could predict ecDNA patterns in another cell line with moderate success, indicating that the framework holds potential for cross-cell-line applications. Similarly, models trained on images of untreated cells were able to predict ecDNA behavior in drug-treated cells with reasonable accuracy, demonstrating that the training process is sufficiently robust to generalize across treatment conditions. These findings suggest that the system is flexible and capable of adapting to varying biological contexts, though further validation will be needed to fully assess the extent of this generalizability.

## Methods

### Primary Cell Culture

NCIH2170, SNU16, and NCIH716 cells were purchased from ATCC and were grown in RPMI media (Gibco) supplemented with 10% heat-inactivated fetal bovine serum (Gibco) in a humidified incubator with 5% CO_2_. SUM159PT cells were given as a gift from Gary Johnson’s laboratory (UNC Chapel Hill) and were STR verified and checked for mycoplasma. SUM159PT cells were grown in Ham’s F-12 media supplemented with 5% heat inactivated fetal bovine serum along with 10mM HEPES, 1ug/ml Hydrocortisone, and 5ug/ml Insulin. Cells were harvested with 0.25% Trypsin in DPBS (Gibco). Viable cells were counted using a Countess 3 (Invitrogen) counter and trypan blue (Invitrogen, T10282). Cells were collected for metaphase and karyotype (G-banding) experiments within 3 passages.

### Metaphase Analyses

Cells were arrested at metaphase by overnight colcemid treatment (12-20h) at 0.1 µg/mL (10 µg/mL Colcemid Solution, FUJIFILM Irvine Scientific) in cell culture media, when cells were ∼70% confluent. Cells were harvested per standard cell culture procedure with minor alterations that trypsinization is quenched by a mixture of cold colcemid-spiked media and PBS wash, to maximize the yield for semi-/adherent cells. Harvested cells were resuspended in 1 mL of 1x PBS by pipetting and transferred to a 1.5 mL microcentrifuge tube for centrifugation at 5000 rpm for 2 minutes. The supernatant was aspirated and each pellet was incubated with 600 µL of pre-warmed 37°C 0.075M KCl (Gibco) and at 37°C for 15 minutes in a water/bead bath. To quench the reaction we prepare Carony’s fixative (3:1 methanol:glacial acetic acid) fresh and add it dropwise in the tube. Tubes were immediately centrifuged at 5000 rpm for 2 minutes. We left 100-200 µL of supernatant to suspend pellets followed by addition of 600 µL of fixative dropwise and resuspension. This step was repeated 3 times. Depending on cell counts, we added 0-1 mL of fixative dropwise to make a 5-6 million cells/mL suspension. This ensures the optimal cell density for discernible spacing of single-cell karyotypes when dropped and imaged on microscope slides.

### Image Generation

Generating condensed chromatin during metaphase allows for the best imaging of ecDNA. Before karyotyping, cells undergo a 4 stage preparation, are arrested at metaphase, are incubated in a hypotonic solution, undergo cell fixation and staining.

Using cells cultured by the Brunk Lab, samples are first introduced to a colcemid treatment of 0.1 µg/mL in cell culture media once around 70% confluency is achieved. This treatment arrests cellular division during mitosis by binding to tubulin preventing spindle formation and cytokinesis. Cells are then collected with standard culture procedure then resuspended in 1mL of 1xPBS via pipette. Centrifugation of the suspension in 1.5mL tubes takes place at 5000 rpm for 2 minutes. In the next stage, the cells are first incubated with 37°C 0.074M KCl, which is added dropwise with slight agitation to resuspend the cells. After 15 minutes at 37°C the cells will have swollen and are fragile through osmotic pressure and ready for fixation. For each sample, 600 µL of Carony’s fixative (3:1 methanol to glacial acetic acid) was transferred dropwise before another 2 minutes in the centrifuge. We remove all but about 150 µL of supernatant to then agitate the remaining fixed cells until they are resuspended. Another 600 µL of fixative was transferred, agitated, then centrifuged for 2 minutes at 5000 rpm. This fixation process was repeated three times, but on the last cycle, the amount of fixative added was only enough to make ∼6 million cell/mL suspension (0-1mL).

With cells at the adequate density for single cell imaging, a drop of the solution was dropped upon a humidified microscope slide from an arms length above the slide. Cells on the slides were left to dry for an hour. They were equilibrated in 2X saline-sodium citrate then dehydrated using an increasing gradient (70%, 85%, 100%) of alcohol solutions for 2 minutes each. Slips were then stored at 37°C for 16-20 hours using a slide moat. Slides were then washed in 0.4X saline-sodium citrate and again in 2X saline-sodium citrate 0.05% Tween20 for 2 minutes. After a final dip in 2X saline-sodium citrate, a drop of SlowFade Diamond Antifade Mountant with DAPI was added to the center. The slides were put in another coverslip and sealed with nail polish then used in imaging.

### Image ranking

Images were evaluated on a 0 to 4 scale, with 0 indicating the lowest confidence in image quality and 4 representing the highest. The ranking process considered factors such as image resolution, clarity, and human oversight. The initial assessment focused on the content and visual quality. Resolution was reviewed to identify blurriness, out-of-focus chromosomes, or artifacts. This preliminary review set a ceiling for the final score; for instance, if the resolution was insufficient to clearly resolve individual chromosomes or ecDNA, the image could receive a maximum score between 0 and 2. In contrast, images with sharp features that clearly resolved chromosomes and ecDNA could achieve a higher ranking of 3 or 4.

Another factor considered was the distribution of material around nuclei, referred to here as the nuclear splash pattern. In some cases, nuclear material spreads unevenly, influenced by factors like nearby artifacts or popped nuclei, which disrupt the typical elliptical shape of a nucleus. The rank was reduced only if these artifacts caused significant overlap between regions, making it difficult to confidently distinguish boundaries. Images with highly crowded fields or poorly defined ROI boundaries were also downgraded, as such conditions made consistent annotation across different users more challenging.

After evaluating image quality, the annotations were reviewed for accuracy. Merged images with multiple probe layers were separated to isolate the DAPI stain, and the original annotations were re-imported. To enhance visibility of small ecDNA, the contrast was increased. Annotations were then verified based on their proximity to the original markings. If most annotations aligned within a few pixels of the original, the image was assigned a rank of 4. However, the rank was lowered if significant errors were detected. False positives were noted when annotations overlapped with chromosomes, debris clouds containing probe molecules, or when ecDNA was duplicated due to layer misalignment or indicator splitting. False negatives occurred when dim ecDNA was missed, when ecDNA was too close to chromosomes or artifacts, or when clusters of ecDNA were undercounted. If such errors were prevalent, the image was downgraded to a rank of 1. In total, we had 183 images marked as 0, 290 images marked as 1, 700 images marked as 2, 970 images marked as 3, and 441 images marked as 4.

Finally, each image was assigned a final rank, and detailed notes were recorded. These notes included suggestions for improving annotations to increase the image’s rank, as well as any observations—such as uneven brightness or scaling—that could explain deviations in accuracy across images but were not directly captured by the ranking criteria.

### ROI segmentation approach

Images were isolated to the single-nucleus level by cytogeneticists, technicians, or scientists who defined the perimeter of the region of interest. This was accomplished using the lasso tool in the FIJI software(Schindelin et al. 2012, Schneider et al. 2012) to create a binary image representing the region of interest (ROI) for each nucleus. Masks were then generated for each image to tightly constrain the focus to the annotated regions and chromosomes. This step aimed to minimize the inclusion of unannotated data that could interfere with model training. However, in some cases, artifacts such as unpopped nuclei could not be fully excluded when defining the ROI. These artifacts were allowed to remain in the final ROI to provide counterexamples for the model, which could enhance its robustness.

### Expanding pixels around annotations

During the original annotation phase, the FIJI counting tool was used by multiple different operators/technicians/scientists to document the count. This tool will give a total count of the annotations made to the image and tag a single pixel the location. Each image was 2048 by 2448 pixels and could have up to a thousand annotations. Due to the large class imbalance each annotation was expanded using a python script to a 7 pixel tall 7 pixel wide diamond patch to decrease the class imbalance and build in padding for each annotation not being exactly in the center of each ecDNA. The pixel patch was transferred as a perimeter into a single .npz file that would translate to a segmentation by MIA.

### Performing Analysis with MIA

Within MIA, each image was dissected into 224 × 224 pixel patches and randomly augmented with any affine transformation. The patches were then fed into a U-Net with a Res-Net-50 encoder neural network. The model used a focal conservative loss with a class weighing of 0.82. The batch size (number of images per model update) was set at 8, and the learning rate was set 0.001 with a 0.1 reduction factor for loss plateaus over three epochs until 1.00 * (10^−6^). Contours were not predicted during the training due to memory limitations, but were optimized to match model training data ecDNA counts in post-processing. These hyperparameters were tuned based on the minimum loss and computational resources over 50 epochs of a reduced dataset.

For training of large models, the model was trained for between 65 and 75 epochs based on the final plateau of loss. Contours were separated in the masks to segregate overlapping ecDNA based on ecDNA count. The segmentation masks were then exported from the .npz coordinates into a .tif image.

A detailed guide on how to prepare, rank, and analyze images using the MIA with FISH images is provided in the Supplementary Information. This documentation includes step-by-step instructions on image preparation, ranking criteria, and running MIA for optimal results.

### Accessing Accuracy of Predictions

To assess the model’s performance, we used a standard accuracy metric defined as the difference between the total predicted count and the total ground truth count, normalized by the ground truth total. This metric provides a measure of the model’s deviation from the true count. The majority of the observed error resulted from the model underestimating the true count, indicating a tendency toward undercounting.

### Error Metric and Model Evaluation

To evaluate the model’s performance, we calculated the Mean Absolute Error (MAE) between the ground truth ecDNA count and the predicted ecDNA count. The MAE is defined as:

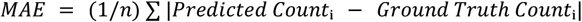

This metric provides an average of the absolute differences between predicted and actual counts, offering a straightforward measure of the model’s accuracy. We selected the MAE because it reflects the magnitude of the prediction error without being influenced by the direction of the error (over- or underestimation).

### Precision

Two measures of precision were calculated: location-based precision and count-based precision. Location-Based Precision

This metric evaluated precision at the pixel level by determining whether individual pixels from the model’s segmentation output overlapped with the ground truth map. Each pixel present in both the ground truth pixel patch and MIA’s segmentation output was counted as a true positive. Conversely, pixels present in the model’s output but not in the ground truth were counted as false positives. In this method, each pixel was treated as an individual label, providing a highly granular precision measure.

### Count-Based Precision

This measure compared the overlap between predicted segmentation labels and the corresponding ground truth pixel patches. If at least one pixel of a predicted segment overlapped with the corresponding ground truth patch, the segment was considered a true positive. To avoid overestimation, any predicted segment overlapping multiple ground truth patches was counted only once as a true positive. The total number of overlapping segments was divided by the total ground truth segment count to yield the precision score.

Any difference between the total number of predicted segments and the overlapping true positives was recorded as the false positive count.

### Object-Oriented Model Performance Evaluation

One approach to assess model performance was object-oriented evaluation. In this method, the true positives (TPs) were calculated by counting the number of ground truth (GT) objects that overlapped with at least one predicted object, while subtracting the number of predicted objects that overlapped with more than one GT object. This ensures that each true positive consists of a unique pairing between one predicted object and one GT object, avoiding overcounting.

The false positives (FPs) were determined by subtracting the total number of true positives from the total number of predicted objects. Similarly, the false negatives (FNs) were computed by subtracting the number of true positives from the total number of GT objects.

The true negatives (TNs) were identified by counting the pixels labeled as zero in both the prediction masks and the GT labels. To adjust for the disproportionate number of background pixels compared to ecDNA pixels, we scaled the number of true negatives by dividing it by the ratio of background pixels to ecDNA pixels in the GT for each image. This result was further divided by 25, representing the size of each GT object, to ensure that the number of true negatives was comparable to the number of true positives.

This scaling ensures that the accuracy metric reflects the importance of true positives and is not disproportionately influenced by the abundance of true negatives in the data.

### High-Confidence Image Set and Model Performance

To estimate the highest achievable accuracy, a curated set of 194 high-confidence images was used. These images were stained exclusively with a DAPI probe, and their annotations were carefully reviewed and corrected. The regions of interest (ROIs) were meticulously defined to minimize noise and extraneous content while ensuring that all annotated ecDNA remained in view. Special attention was given to the ROI edges to avoid abrupt cuts into the bright fields surrounding objects. Any images with artifacts or blur within the ROI were excluded from the dataset.

The dataset was split into a training set of 131 images and a test set of 63 images, selected randomly. The model was trained using the same procedures and parameters as the main model (ensure that this is referenced if detailed elsewhere), for a total of 60 epochs. Model performance on the validation set resulted in a mean absolute error (MAE) of 6.84%, while the training set predictions achieved a MAE of 5.14%.

### Estimating User-to-User Variability

Given that the data inherently reflects biases introduced by the human operators (technicians/scientists), we aimed to estimate the extent of user-to-user variability. To do this, we conducted a test with 11 images, where multiple operators independently annotated and counted the same images in both the DAPI-only layer and the merged images.

The analysis revealed an average error of approximately 8.4% for the DAPI-only images, while the merged images exhibited a higher variance, with an average error of 12.0% between users.

## Data availability

The workflow for this project has been compiled into a series of user-friendly Jupyter notebooks available at: https://github.com/Brunk-Lab/ecIMAGE.

## Acknowledgements

The authors gratefully acknowledge the funding support provided by the IBM Junior Faculty Development Award from the University of North Carolina at Chapel Hill (to EB), the Computational Medicine Pilot Award (to EB) and the NIGMS Grant 2R25GM089569-14 (to DG). We also extend our appreciation to the Renaissance Computing Institute (RENCI) at the University of North Carolina at Chapel Hill, with special thanks to Dr. Matt Suttusky, Prof. Ashok Krishnamurthy for their valuable insights and time spent discussing key concepts and theories. We also appreciate the discussions with Prof. Marc Niethammer and Qin Lu on computer vision.

## Author Contributions

EB conceived and managed the research. EB led, designed, conducted analyses, oversaw experiments and manual annotations. KG and AM conducted computational analyses. DG, JF, WN, and DC conducted additional computational analyses. WD, OC, NG and JC conducted FISH experiments. KG, AM, WD, OC, JC, NG, YW, KS, JF conducted manual counting. JC trained WD and OC to perform FISH experiments and all students to manually count images. EB wrote the manuscript. EB, KG and AM created figures and tables for the manuscript. All authors read and approved of the manuscript.

## Notes

https://github.com/Brunk-Lab/ecIMAGE.

### Competing Interest Statement

The authors have declared no competing interest.

